# Habitat use of juvenile green sea turtles along the North Pacific coast of Costa Rica

**DOI:** 10.1101/2025.06.02.657549

**Authors:** Fanqi Wu, Veronica Valverde-Cantillo, Chelsea Durr, Mario Espinoza, Maike Heidenmeyer, Christopher G. Lowe, James R. Spotila, Frank V. Paladino

**Affiliations:** Department of Biological Sciences, Purdue University Fort Wayne, Fort Wayne, Indiana, United States of America; The Leatherback Trust, Fort Wayne, Indiana, United States of America; Global Cause Foundation, Blacksburg, Virginia, United States of America; Centro de Investigación en Ciencias del Mar y Limnología, Universidad de Costa Rica, San José, Costa Rica; MigraMar, Bodega Bay, California, United States of America; Equipo Tora Carey, El Jobo, La Cruz, Guanacaste, Costa Rica; Department of Biological Sciences, California State University, Long Beach, California, United States of America; Department of Biodiversity, Earth and Environmental Science, Drexel University, Philadelphia, United States of America

## Abstract

Understanding how threatened marine species use coastal areas and the extent of connectivity across different spatial and temporal scales is important for identifying critical habitats that can enhance conservation efforts in other regions of their distribution. In this study, we investigated the site fidelity and habitat use of juvenile yellow morphotype green sea turtles (*Chelonia mydas*) in the Gulf of Santa Elena, north Pacific coast of Costa Rica. Fifteen juvenile green sea turtles (49 – 83 cm curved carapace length; CCL) were monitored for 19 – 629 days with an array of 11 acoustic receivers placed within 5 main habitat types: muddy areas, reef patches, macroalgae, rocky reefs, and mangroves. Daily and seasonal locations of the turtles were estimated using a positioning estimation algorithm to determine the site fidelity and habitat use within the study area. Large juveniles (≥ 65 CCL) were detected disproportionally in the macroalgae habitat during the upwelling season (43.9% detections) from December – April, and in the reef patch habitat during the non-upwelling season (36.0%) from May to November. Small juveniles (< 65 cm CCL) had more detections in the reef patch habitat during both seasons (Upwelling: 35.4%, non-upwelling: 45.4%) relative to other habitats sampled. More than half of the turtles monitored (67%) showed strong site fidelity to the study area, ranging from 68.9 to 100% from October 2020 to September 2022. Our results suggest that juvenile green sea turtles, particularly larger individuals, have a strong association with coral / rocky reef patches and macroalgae habitats, similar to adults. Our finding suggested that protecting similar habitats in other areas along the Central American coast will help rebuild the Eastern Pacific Ocean green sea turtle population.

## Introduction

Knowledge of how threatened species move and use specific areas is essential to assess the level of population connectivity and viability. This information can also help identify critical habitats (e.g., feeding and reproductive grounds) for the survival of a species and guide spatial conservation efforts to rebuild their population [1]. The green sea turtle (*Chelonia mydas)* is an endangered species with a declining population [2]. In the Pacific Ocean, the green sea turtle population is divided into two subgroups that can be recognized based on coloration and shape: the yellow morphotype that breeds in the western Pacific (yellow sea turtle, or green sea turtle) and the black morphotype that breeds in the eastern Pacific (black sea turtle) [3–5]. The main morphometric differences between the two subgroups are the shape of the head, carapace, plastron, and flippers [5]. Both morphotypes spend their early pelagic stage in the open ocean until they grow to about 44 cm curved carapace length (CCL), and then they settle in coastal foraging grounds as juveniles for their neritic stage [6].

During the pelagic stage, green turtles are primarily omnivorous and feed on planktonic algae, crustaceans, ctenophores, and jellyfish [7]. Once juvenile green sea turtles move to coastal foraging grounds, turtles undergo a shift in feeding behavior from an omnivorous diet to a predominantly herbivorous one, feeding on macroalgae, seagrass, and/or mangrove material [7–9]. Here, coastal juveniles and sub-adults use a variety of habitats including sand, seagrass beds, reef flats, terraces, and mangrove areas [10,11]. For example, in the northern Gulf of Mexico, juveniles use sand and seagrass areas [10] while in Kaneohe Bay, Hawaii, juvenile green sea turtles use reef patches or coral-covered areas for resting and foraging (Brill, et al., 1995). Often, these coastal juvenile foraging habitats persist into adulthood and become the adult foraging habitat used between nesting seasons, therefore understanding which habitats green sea turtles select within the given options and how they use these options is important to long-term conservation. Further, the bays along the Pacific coast of Central America may be important developmental habitats for juvenile green turtles and there is currently limited knowledge about these populations [12].

In the eastern Pacific region (EPO), habitat use spatial studies must address the seasonality that exists here. While some populations of juvenile green sea turtles demonstrate fidelity to a single foraging habitat, others shuttle between habitats in response to water temperature changes, habitat quality changes, or prevailing currents [10,13–15]. Because of the multi-decadal El Niño Southern Oscillation (ENSO), seasonal upwelling, and the resulting fluctuations in nutrient levels, plankton populations, and other factors that influence the entire food web [16], we expect habitat selection, foraging ecology, and space use of juvenile green sea turtles to be equally seasonal and dynamic [17]. To address this, we used passive acoustic telemetry [18] to determine site fidelity and habitat use of juvenile yellow morphotype green sea turtles on the north Pacific coast of Costa Rica. Our study was conducted in Matapalito and Santa Elena Bays in the Gulf of Santa Elena as these two bays provide a unique habitat combination of macroalgae, reef patches, rocky reefs, muddy areas, and mangroves. Both morphotypes of green sea turtles inhabit this area and stable isotope analysis indicates that yellow morphotype green sea turtles mostly feed on macroalgae [17,19]. By using acoustic telemetry and satellite imagery across natural seasonal variability, we can identify critical habitats for green sea turtles and monitor turtles’ response to the water temperature and nutrient changes. These data provide important information not just about how sea turtles use their available habitat today but may help us predict how they will adapt their habitat use to future scenarios.

## Methods

### Ethics Statement

This study was carried out in accordance with institutional, federal, and international guidelines and permits. All data were collected under the protocol approved by the Purdue Animal Care and Use Committee (protocol #1804001737). Permissions to work with endangered sea turtles in Costa Rica were granted under permits from Guanacaste Conservation Areas (ACG) of the Ministry of Environment of Costa Rica (ACT-OR-DR-099-2022, ACT-OR-DR-120-2022).

### Study Area

Santa Elena Bay (SEB) (10°55’10.39"N, 85°47’52.88" W) and Matapalito Bay (MPB) (10°55’58.75"N, 85°47’29.48"W) are in the Gulf of Santa Elena, on the north Pacific coast of Costa Rica. SEB is a relatively large bay (7.2 km^2^) with a wide variety of habitats, ranging from mangroves, soft-mud, and gravel to rocky/coral reefs. MPB is a small bay (0.72 km^2^) adjacent to SEB, with rocky and coral reef patches, macroalgae, and sandy substrate. Both SEB and MPB are influenced by the strong northeast trade winds and seasonal upwelling. The upwelling season is from December to April, and the non-upwelling season is from May to November [20,21]. During the upwelling season, shallow coastal waters are cold (∼20 °C) and nutrient-rich, and typically have low pH and low oxygen levels [22], which results in rapid growth of macroalgae [23]. During the non-upwelling season, the water temperature is warm (∼28 °C), and the area has less macroalgae [24].

### Acoustic Receiver Placement

We used seven VR2Tx and four VR2W acoustic receivers (InnovaSea Systems Inc., Nova Scotia, Canada) to record detections from animals tagged with acoustic transmitters. The detection range was variable and influenced by the habitat, with recorded ranges of less than 200 m in rocky and reef habitats to up to 600 m in estuarine areas from SEB (V. Valverde unpublished data). Hourly water temperature was measured by the VR2Tx receivers (± 0.5 °C) or HOBO Water Temperature Pro v2 Data Loggers (Onset Computer Corporation) (± 0.2 °C) attached to the VR2W receivers. Prior to installation, we covered each receiver with electric tape and a nylon stocking to prevent biofouling. Animals attaching directly to the receiver can reduce detection efficiencies (Heupel et al. 2008). For receiver placement, we selected areas with macroalgae, reef patches, mangroves, soft-mud bottoms, coral reefs, and rocky reefs in both bays (Fig 1). Receivers were supported on stainless steel bars or attached directly to a mooring line using cable ties and placed directly on the substrates. In SEB, 7 receivers (3 VR2Tx and 4 VR2W) covered mangrove, muddy, and rocky reefs habitat types. In MPB, 4 VR2Tx receivers covered reef patches, macroalgae, and rocky reefs habitat types (Fig. 1.). Each receiver recorded the date, time, and a unique ID code whenever it detected an animal tagged with a transmitter. We manually downloaded data from the receivers every 2–3 months.

**Fig 1.**
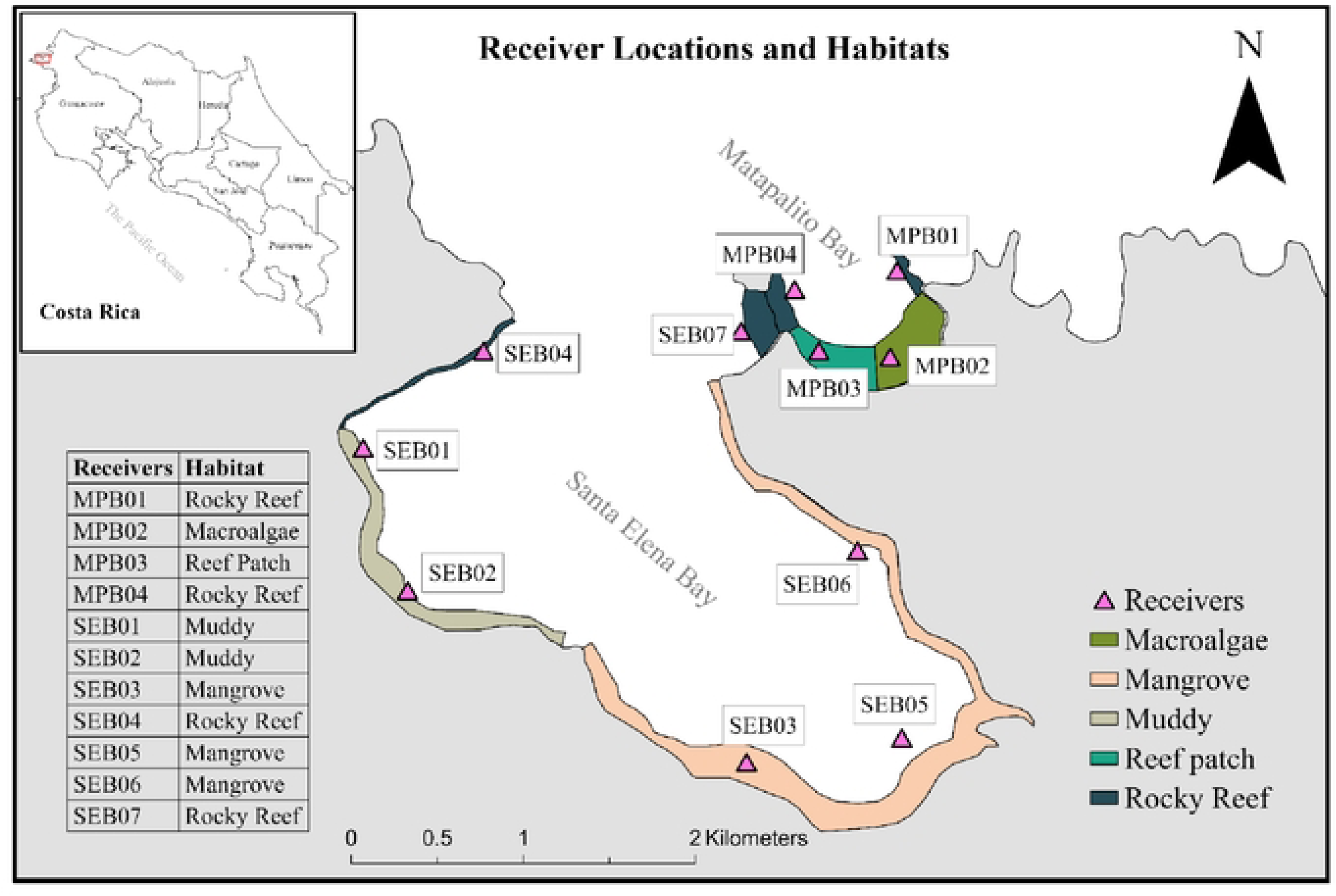
Receiver locations and habitats. Seven receivers were placed in SEB and 4 receivers were placed in MPB along the north Pacific coast of Costa Rica. MP02 is coarse sand and macroalgae, but the macroalgae is seasonal. SEB 3 and SEB 5 habitat is soft mud. SEB 6 is coarse gravel and mangrove with some patchy rocky reef.

### Turtle Capture

We captured 15 green sea turtles in MPB in cooperation with field assistants from the Equipo Tora Carey (ETC) monitoring program. We used two methods to catch turtles: turtle tangle nets and by hand. Tangle nets consisted of three nets attached together (240 m long by 8 m deep) with a mesh size of 42 cm. The top of the net was held up with small buoys at 2 m intervals, and the bottom of the net was weighed down with small weights. If a turtle was caught in the nets, the net was light enough for the turtle to drag the whole net to the surface to breathe [17]. We checked the nets every 30 min or less as well as when something appeared to be in the net. The nets were placed in the morning adjacent to the reef patch area near receiver MPB03, where the water was about 11 m deep [17]. After placing the nets in the water, the ETC crew snorkeled in the shallow water areas nearby to look for turtles to hand capture.

Once caught, turtles were transferred to the research boat for morphometric data collection. First, we identified each turtle with either an INCONEL metal identification tag (Style 681, National Band and Tag Company USFWS tags) or a PIT (Passive Integrated Transponder) tag (FriendChip, Avid Identification Systems, Inc.). Next, we measured CCL and curved carapace width (CCW) using a soft measuring tape (± 0.5 cm). We measured CCL from the anterior point at the midline (nuchal scute) to the posterior tip of the supracaudal scute. We measured CCW across the widest part of the carapace. Finally, we measured the body mass of each turtle using a Rapaura 50 kg spring scale (± 0.1 kg) and a heavy-duty nylon rope.

### Acoustic Transmitter Installation

We installed VEMCO v16 coded acoustic transmitters (InnovaSea Systems Inc., Nova Scotia, Canada) on each turtle. Each transmitter had a length of 68 mm and a mass of 10.3 g in water and sent a random signal pulse every 60–90 seconds. We installed each transmitter on a clean and flat area of the marginal scute just above the left or right rear flipper [25]. We used a wire brush to clean the carapace and an alcohol pad to sanitize the area then used a 5/16” drill bit to drill 3 holes in the marginal scutes on the posterior carapace. We used stainless steel wire and cable crimp sleeves to tie the transmitter to the carapace and coated the base of the transmitter in marine glue. It took about an hour for the glue to dry, during which time the turtle was monitored on the boat. Once the transmitter was secured and the glue was dry, we released the turtle back into the water.

### Data Analysis

This study mainly focused on habitat use but we also looked at site fidelity and estimated each turtle’s home range and core use area. The habitat use of each turtle was determined by its hourly location data [26]. We categorized every time a receiver recorded the signal from a transmitter as a detection. If a turtle stayed in a receiver detection range for 60 min without moving to another receiver, we counted it as one detection. If a turtle visited multiple receivers in 60 min, we counted each as a detection.

Site fidelity was calculated as the number of days an individual was detected in the two bays relative to the number of days of monitoring period and expressed as a percent. For each turtle, the tracking period was defined as the number of days between the first and last date detected, and the detection days were defined as the number of days an individual was detected over the tracking period (Table 1). Given that turtles larger than 65 cm CCL are known to exhibit similar feeding behaviors to adults and may be found in similar habitats [19,27,28], we separated the individuals into large (≥ 65 cm CCL) and small (< 65 cm CCL) size categories [9]. Turtle CM01 was only detected for 4 days, so we did not include it in subsequent analyses.

**Table 1.** Turtle and Tracking Information. 15 yellow morphotype green sea turtles (6 large turtles and 9 small turtles) were tracked in Matapalito and Santa Elena bays in northwest Costa Rica.

| **Turtle IDs** | **Start Date** | **CCL (cm)** | **CCW (cm)** | **Mass (kg)** | **Tracking Period** | **Detection Days** | **Size Group** |
| --- | --- | --- | --- | --- | --- | --- | --- |
| CM01 | 10/16/20 | 56.0 | 53.0 | 18.0 | 19 | 4 | Small |
| CM02 | 12/06/20 | 55.0 | 52.5 | 19.1 | 628 | 185 | Small |
| CM03 | 12/06/20 | 53.5 | 47.0 | 18.0 | 629 | 258 | Small |
| CM04 | 01/27/21 | 65.0 | 62.5 | 34.5 | 134 | 121 | Large |
| CM05 | 03/16/21 | 66.0 | 59.0 | 31.0 | 396 | 171 | Large |
| CM06 | 06/16/21 | 83.0 | 70.0 | n/a | 363 | 82 | Large |
| CM07 | 08/21/21 | 53.3 | 48.6 | 18.7 | 379 | 261 | Small |
| CM08 | 10/11/21 | 49.8 | 46.8 | 15.5 | 327 | 299 | Small |
| CM09 | 10/11/21 | 53.3 | 48.5 | 16.7 | 326 | 264 | Small |
| CM10 | 10/11/21 | 54.3 | 48.6 | 20.0 | 301 | 300 | Small |
| CM11 | 10/11/21 | 65.0 | 59.6 | 28.0 | 310 | 282 | Large |
| CM12 | 03/26/22 | 49.0 | 48.5 | 12.8 | 162 | 162 | Small |
| CM13 | 04/28/22 | 71.5 | 66.0 | 45.5 | 128 | 126 | Large |
| CM14 | 04/28/22 | 65.0 | 60.9 | 32.7 | 128 | 114 | Large |
| CM15 | 07/15/22 | 50.5 | 49.5 | 14.0 | 51 | 51 | Small |

We used the Minimum Bounding Geometry (MBG) tool in ArcGIS Pro (v2.9, ESRI 2022) to calculate the activity area size [10,13]. Since the acoustic receivers and transmitters only provided locations of the receivers, we used the Count Overlapping tool in ArcGIS Pro to calculate the core use area used by all 14 turtles.

We identified each habitat area by combining satellite images (Google Earth Pro 7.3.6) from different seasons and field observations and digitized the area of each habitat type. Field observations were conducted using snorkel transects in Matapalito Bay. We quantified vegetation for each season individually. Then we calculated the size of the combined areas for each habitat type. We used Spearman’s Correlation in RStudio (R Studio Inc., Boston, MA, USA) to analyze the correlation between CCL and activity areas.

## Results

### Activity Area and Site Fidelity

Fifteen juvenile yellow morphotype green sea turtles (mean CCL: 59.6 ± 9.9 cm) were tracked for 19 – 629 days. For analysis, we only included 14 turtles (51 to 629 d). Individuals were detected between 51 and 300 days. CM15 had the shortest tracking period (51 days) and was detected on each day. CM02 and CM03 had the longest tracking periods (628 and 629 days) and were detected for 185 and 258 of those days, respectively. CM06 was tracked for 363 days, but it was only detected in the study area for 82 of those days, whereas CM05 was tracked for 396 days and was detected for 171 of those days. Site fidelity of the 14 turtles ranged from 22.6 to 100%. Four turtles (CM02, CM03, CM05, and CM06) had low site fidelity (22.6 – 43.2%), while the remaining 10 individuals had high site fidelity (68.9 – 100.0%). Six of the 14 turtles were detected only in MPB (CM05, CM07, CM10, CM11, CM12, and CM15); therefore, their activity areas were considerably smaller (0.1 – 0.3 km^2^). CM09 was detected by all the receivers and CM14 was detected by all receivers except SEB03.

The estimated activity area for all turtles ranged from 0.1 to 5.4 km² (mean ± SD: 1.73 ± 1.91 km²). The core use area was 0.13 km² in MPB. This core area covered a reef patch and macroalgae habitats and was used by more than 12 turtles. There was a moderate positive relationship between CCL and activity area, but it was not statistically significant (S = 241.92, rho = 0.47, p = 0.09).

### Habitat Use

Turtles were detected disproportionately in the reef patch, rocky reef, and macroalgae areas in comparison to the proportion of the study site that these habitats occupied. Overall, the detection percentages in each habitat type during the study period were 38.6% in reef patch, 27.4% in rocky reef, 20.6% in macroalgae area, 12.4% in muddy area, and 1.0 % in mangrove area. The estimated area size of all habitat types in both bays was 1.19 km^2^, of which 53.2% was mangroves, 15.5% was sandy areas, 11.6% was macroalgae areas, 11.4% was rocky reef areas, and 8.2% was reef patch areas (Table 2).

**Table 2.**
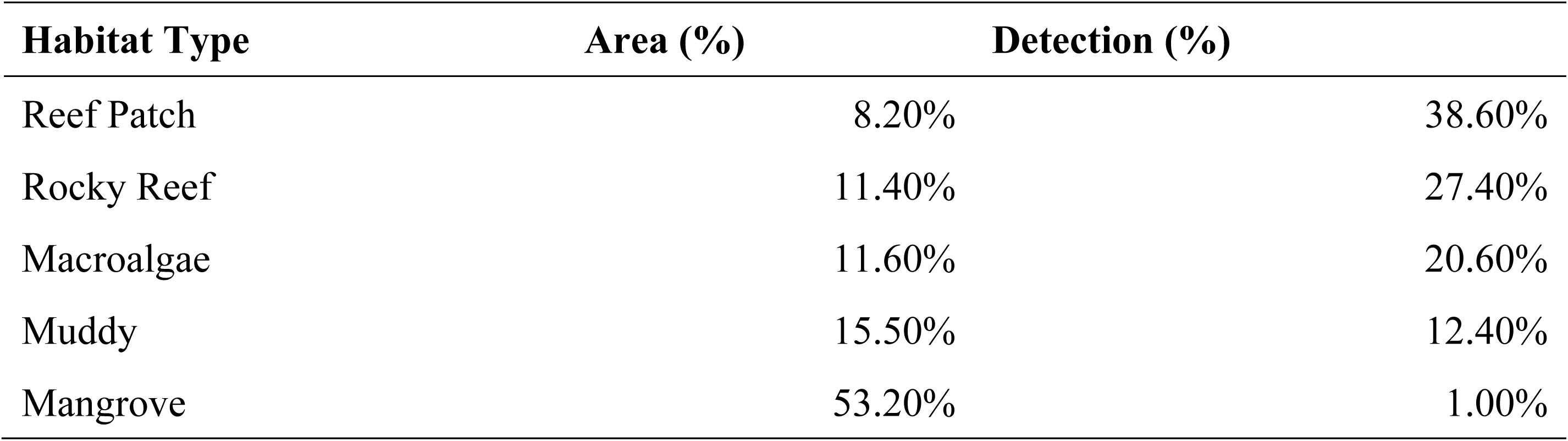
Comparison of the percentage of areas of each habitat and the detection of yellow morphotype green turtles in those habitats in Matapalito Bay and Santa Elena Bay in northwestern Costa Rica.

Large and small turtles used habitats differently. Over the entire study period, most detections of large turtles were from the reef patch area (34.2%). Large turtles had 30.2 % of their detections from the macroalgae area and 17.0% from the rocky reef area in MPB (Fig 2a). Most detections of small turtles were from the reef patch area (41.1%), followed by 16.7 % of detections in the rocky reef area in MPB, 15.1% from the macroalgae area, and 14.1% from the muddy area in SEB (Fig 2b).

**Fig 2.**
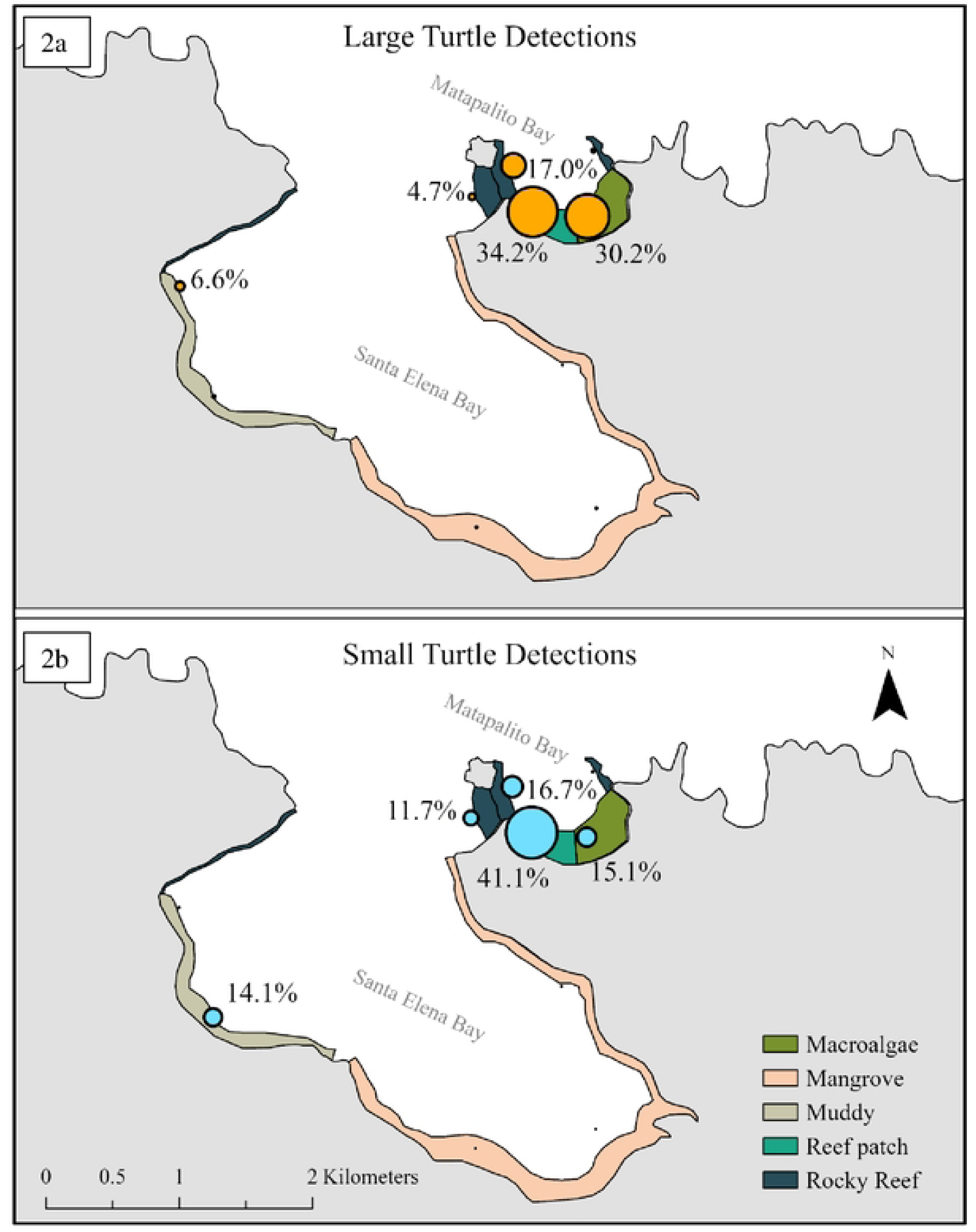
Habitat Use by Size Classes. Detection percentages of large (a) and small (b) juvenile yellow morphotype green sea turtles in different habitats in Matapalito Bay and Santa Elena Bay in northwestern Costa Rica Sea water temperatures ranged from 15.0 to 40.0 °C. Monthly mean water temperatures ranged from 20.3 to 29.2 °C (mean ± SD: 26.5 ± 2.2 °C) during the study period. In the upwelling season, monthly mean water temperature was from 21.4 °C to 25.6 °C (23.8 ± 1.4 °C) in MPB and from 20.3 to 27.4 °C (24.6 ± 1.8 °C) in SEB. In the non-upwelling season, monthly mean water temperature was from 26.3 to 29.2 °C (27.8 ± 0.7 °C) in MPB and from 26.1 to 29.2 °C (28.1 ± 0.7 °C) in SEB.

During the upwelling season, most detections of large turtles were from the macroalgae area (43.9%). Large turtles had 32.4% of their detections from the reef patch area and 20.0% from rocky reef area in MPB. No large turtles were detected in SEB, except from rocky reef areas (0.2%). For the small turtles, most detections in the upwelling season were from the reef patch area (35.4 %). Small turtles had 26.5% of their detections from the rocky reef area in MPB, 19.2% from the macroalgae area, and 13.4% from the muddy area in SEB (Fig 3a). During the non-upwelling season, most detections of large turtles were from the reef patch area (36.0%) (Fig 3b). Large turtles had 17.6% of their detections from the macroalgae area, 14.2% from rocky reef area in MPB, and 12.8% from the muddy area in SEB. During the non-upwelling season, most of detections of small turtles were from the reef patch area (45.4%). Small turtles had 18.2% of their detections from rocky reef area in SEB, 14.6% from muddy area in SEB, and 12.1% from the macroalgae area (Fig 3b).

**Fig 3.**
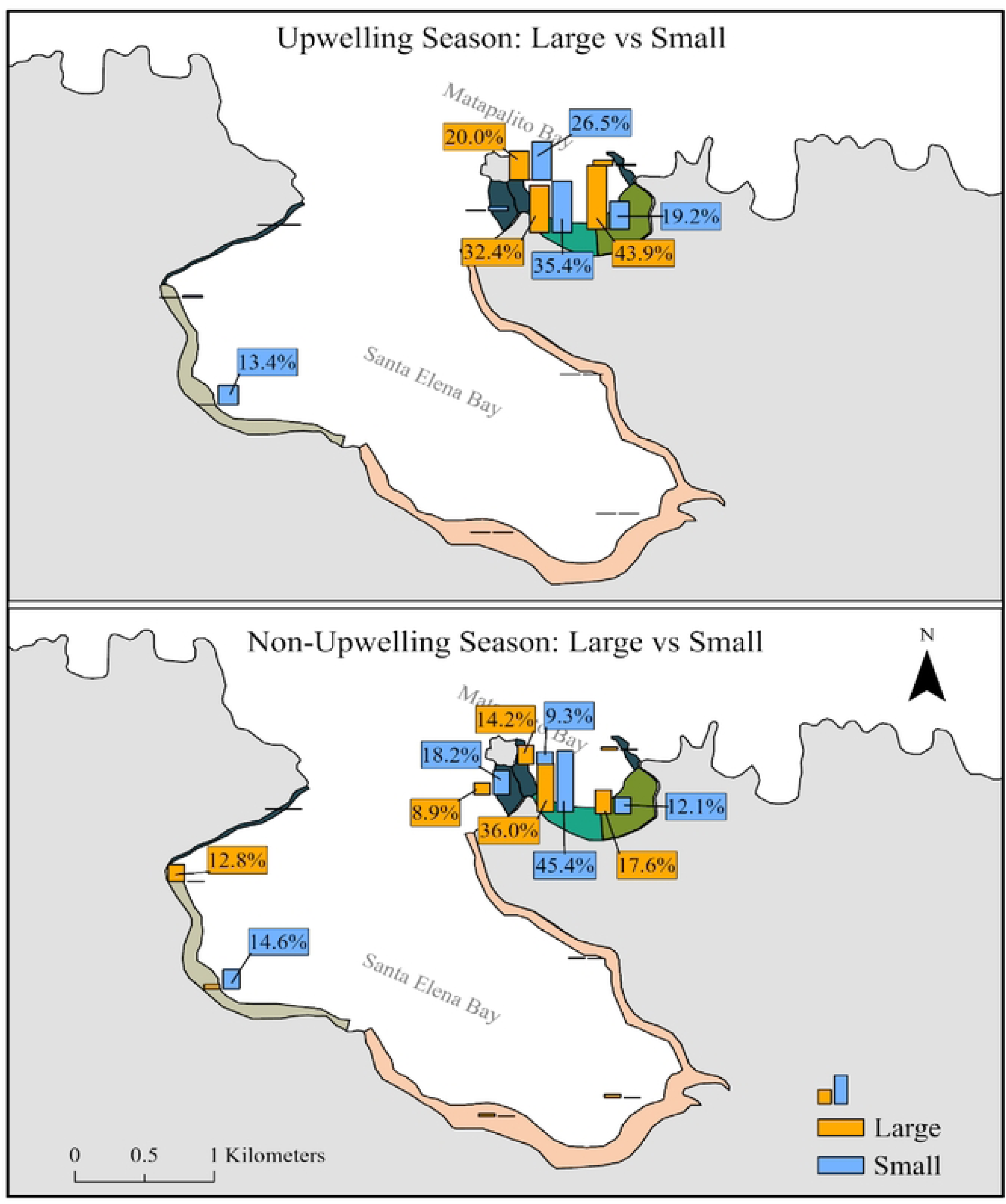
Seasonal Habitat Use. Detection of the juvenile yellow morphotype green sea turtles in Matapalito Bay and Santa Elena Bay in northwestern Costa Rica during non-upwelling season (a) and dry season (b).

During the non-upwelling season, 39.1% of the detections were from the reef patch area and 13.2 % were from the macroalgae area in MPB. During the upwelling season, 31.5 % of the detections were from the reef patch area and 26.7 % were from the macroalgae area in MPB. During the upwelling season, there were more detections of turtles in the rocky reef area near MPB04 (22.2%) than during the non-upwelling season (10.4%). There was a greater use of the area between MPB and SEB (13.9% of detections) during the non-upwelling season than during the upwelling season (1.7%). There was a small difference in the use of the sandy area near SEB02 in SEB during the two seasons.

## Discussion

Our results indicated that yellow morphotype green sea turtles had high site fidelity to the study area, especially to MPB where they were initially captured. However, some turtles spent time away from the study area indicating that they had a larger home range than we could detect. Turtles were detected disproportionately in the reef patch, rocky reef, and macroalgae areas in comparison to the proportion of the study site that these habitats occupied. Large and small turtles used habitats differently. In addition, turtles used habitats differently during the upwelling and non-upwelling seasons.

### Activity Area and Site Fidelity

Our results showed that juvenile yellow morphotype green sea turtles exhibited strong site fidelity to MPB and SEB, especially to MPB where they were initially captured. They did this by either remaining in the bay or making short trips out of the bay and returning. All these behaviors are similar to findings reported in other studies in the Atlantic Ocean [10,29,30], the South-West Indian Ocean [11], and Puerto Rico. In the latter, large juvenile green turtles leave their normal activity areas on brief trips [31]. Our results indicated that those juvenile turtles relied on the food resources in this area. Also, the coral reefs and rocks in this area may provide shelter for juvenile turtles. Food resources and safety play critical roles in juvenile green turtle habitats.

The average estimated activity area for the 14 juvenile yellow turtles was 1.73 km^2^, similar to juvenile green turtles in Northwestern Australia (1.63 km^2^) [32] and for juvenile green turtles (2.38 km^2^) in Palm Beach FL, USA [13]. It was, however, smaller than the home ranges of juvenile green turtles (16.62 km^2^) in the Gulf of California, Mexico [33], in St Joseph Bay (3.54 km^2^) in the US Gulf of Mexico [10], and in Everglades (1004.9 km^2^) in FL, USA [30]. The activity area sizes in this study were larger than the home ranges (0.21 km^2^) of juvenile green turtles in the Southwestern Indian Ocean [11]. MPB provided a variety of habitats and food sources ranging from macroalgae to organisms on rocky reef patches. The quality and availability of resources in the small bay allowed turtles to maintain small activity areas. They seldom moved into SEB which, although larger, did not have the same availability of food resource areas (Fig 1). Isla Pitahaya (MPB04) had rocky reef patches that provided food resources that the turtles exploited, especially in the dry season.

### Habitat Use

Our findings indicated that the spatial and seasonal availability of food resources played a significant role in determining when and how juvenile green sea turtles used different habitats. Shallow reef patches and heavy macroalgae growth in MPB provided areas for foraging and resting, which was similar to that reported for green sea turtles in Kaneohe Bay, Hawaii [34].

Green sea turtles in our study preferred reef patch, rocky reef, and macroalgae habitat types (Table 2), whereas juvenile green sea turtles in St Joseph Bay in Florida, USA, tended to use more seagrass beds and open sandy areas [10].

There were seasonal effects on habitat use observed, especially for large juvenile turtles. During the upwelling season, nutrients are brought to coastal areas and support macroalgae growth [35]. Shallow reef patches and the seasonal growth of macroalgae in MPB during the upwelling season provided optimal areas for juvenile turtles to forage and rest. It also provided a variety of habitats and food sources ranging from macroalgae to organisms on rocky reef patches. Turtles were located more often in shallow reef patches than in the macroalgae area but used a combination of reef patches, rocky reefs, and macroalgae habitat types.

During the upwelling season, there was a notable increase in the number of detections of large turtles in the macroalgae area, most likely due to the seasonal algal abundance. Large juvenile turtles relied heavily on macroalgae, similar to juvenile green turtles in Bermuda [36]. In St Joseph Bay in Florida, USA juvenile green turtles used seagrass beds and open sandy areas [10]. In contrast, small turtles in our study predominately remained in reef patches and rocky reef areas in both seasons. The reef patch area can provide both food resources and benthic shelters for juvenile turtles [10,11]. Even during the dry season when macroalgae were abundant in MPB, small turtles were detected near the reef patch area (31.5%) more than in the macroalgae area (26.7%). Large juveniles were detected in the macroalgae area more than small turtles, and the small turtles used the reef patch area more than the large turtles.

### Conservation recommendations

To our knowledge, this is the first study to use telemetry to investigate the habitat use of juvenile yellow sea turtles along the Pacific coast of Central America. Our findings reveal that juvenile yellow sea turtles had small activity ranges (1.73 ± 1.91 km²) with a primary presence in reef patch areas, macroalgae areas, and rocky reef areas. This underscores the significance of those habitats in supporting the development and growth of juvenile green sea turtles. Therefore, we suggest that similar habitat areas throughout Pacific Central America have the potential to support populations of juvenile green turtles. We conclude that these habitats should be protected in all countries because without developmental habitats the population cannot survive. Since 2018, fishing and boating have not been allowed in SEB and MPB. Some of the turtles in our study occasionally leave the study area for extended periods, sometimes even months, we recommend that the entire Gulf needs to be protected and managed.

## Acknowledgement

We acknowledge The Global Cause Foundation, The Leatherback Trust, Equipo Tora Carey, The Center for Marine Conservation and Biology at Purdue University Fort Wayne, and the Jack W. Schrey Distinguished Professor Found for supporting and funding the study. We thank the Ecology and Marine Conservation Lab at the University of Costa Rica, California State University Long Beach, and Earthwatch volunteers for their assistance.

